# Comparative and population genomics analyses of eared pheasants inhabiting highly varying altitudes

**DOI:** 10.1101/2024.05.21.595214

**Authors:** Siwen Wu, Kun Wang, Xuehai Ge, Sisi Yuan, Dong-Dong Wu, Changrong Ge, Junjing Jia, Zhengchang Su, Tengfei Dou

**Author notes:** Corresponding author(s): Changrong Ge, Junjing Jia, Zhengchang Su, Tengfei Dou. These authors contributed equally to this work (Siwen Wu, Kun Wang, Xuehai Ge).

## Abstract

Oxygen pressure varies dramatically with altitudes on Earth; however, humans and animals thrive at almost all altitudes. To better understand genetic basis underlying adaptation of closely related species to varying altitudes, we annotated and compared the genome of a white eared pheasant (WT) (*Crossoptilon crossoptilon*) inhabiting high altitudes and the genome of a brown eared pheasant (BR) (*C. mantchuricum*) inhabiting low altitudes. Moreover, we compared genetic variations in populations of WT and BR as well as of blue eared pheasants (BL) (*C. auritum*) inhabiting intermediate altitudes, and identified thousands of selective sweeps in each species. Intriguingly, the unique genes and pseudogenes in the two genomes converge on the same set of altitude adaptation-related pathways of four functional categories as genes in selective sweeps in each species. Thus, these species appear to adapt to highly varying altitudes by diverging selection on the same traits via loss-of-function mutations and fine-tuning genes in common pathways.

## Introduction

Faunas have gone through an evolutionary process to gain physiological feature for the adaptation of highly varying oxygen abundance. Populations of humans^1-3^ and many animal species that are naturally adapted to lowlands thrive in highland environments, such as Tibetan chickens^4-6^, Tibetan pigs^7,8^, Tibetan goats^9^, Tibetan sheep^10,11^, Tibetan mastiffs^12^, Tibet cattle^13,14^, ground tits^15^, and yaks^16^, to name a few. The major challenge faced by these populations of humans and animals in highland environments is the low oxygen pressure-induced stress on their physiological systems^17-19^. The long-term adaptation to low oxygen pressure environments has resulted in genetic changes in these human^1-3,14,17,20-27^ and animal^4-7,10-16,19,28-31^ populations. Adaptive mutations of many genes of humans^2,14,17,18,26,32,33^ and animals^16,19,30,31,33,34^ have been linked to highland environments. Moreover, introgression has been shown as a major mechanism for human^1,3,27^ and domesticated animals^9,10,13^ to adapt to highland environments. Despite these great progresses, the understanding of human and animal adaptation to highly varying altitudes is still limited^33^. Particularly, some closely related animal species can rapidly adapt to highly varying altitudes, thus, it is interesting to understand the genetic basis of their adaptation from low-to intermediate-and high-altitude environments.

*Crossoptilon*, belonging to the Phasianidae family in the Galliformes order, is a rare but important genus endemic in China^35^. There are four species in the *Crossoptilon* genus, including Tibetan eared pheasant (TB) (*C. harmani*), white eared pheasant (WT) (*C. crossoptilon*), blue eared pheasant (BL) (*C. auritum*) and brown eared pheasant (BR) (*C. mantchuricum*)^35,36^. These species are found in coniferous forests, mixed broadleaf-conifer forests and alpine scrubs in various parts of China with very different altitudes ranging from 800 m to 5,000 m^35,37-42^. TB and WT are believed to be conspecifics, diverging about 0.5 million years ago^43^. They sympatrically inhabit montane forest at a high altitude of 3,000-5,000 m^35,36,38,40-42^. TB is distributed in southeastern Tibet and adjacent northern India (3,000-5,000 m)^35,38,41^, while WT is found in Qinghai, Sichuan, Yunnan and Tibet (3,000-4,300 m)^35,40^. BL and BR are closely related, diverging about 0.3 million years ago^43^, but allopatrically inhabit different areas^35,36^. BL is only encountered in the mountains of Qinghai, Gansu, Sichuan and Ningxia, at an intermediate altitude of 2,000-3,000 m^38,44^, while BR is mainly distributed in mountains in Shanxi and Hebei and near Beijing, at an altitude of 800-2,600 m^37,39,42^. Thus, WT, BL and BR are excellent models to study the genetic basis for closely related species to adapt to highly varying altitudes.

Recently, Wang et al.^45^ assembled a BR genome and re-sequenced three BR subpopulations and a BL population. Although they provide new insights into genomic changes during the course of BR’s long-term population decline^45^, genetic basis underlying the adaptation of the BR and BL to low (800-2,600 m) and intermediated altitudes (1500-3000 m), respectively, were not investigated. Moreover, with a small contig N50 (0.11 Mb), this BR genome assembly has limitations as a reference genome to study the *Crossoptilon* species. To fill the gap, we recently sequenced and assembled the genome of a WT individual at chromosome-level with a contig N50 of 19.6 Mb, and re-sequenced 10 WT individuals and a BL individual^46^. Combining our data in WT and BL^46^ with those of Wang et al. in BL and BR^45^, we were well positioned to investigate the genetic bases of these closely related avian species to adapt to highly varying altitudes using comparative and population genomics approaches. This study is based in part upon SW’s dissertation^47^.

## Results

### Annotation of protein-coding genes in the WT and BR genomes

To see whether gene contents are related to adaption of closely related avian species to different altitudes, we first annotated the assembled WT^46^ and BR^45^ genomes using a combination of reference- and RNA-based methods (Materials and Methods). Using the coding DNA sequences (CDSs) of 53 well-represented Aves (Table S1) as the templates and a total of 23 RNA-seq datasets from 20 tissues of WT individuals and three tissues of BR individuals (Materials and Methods), we annotated 16,377 and 15,410 protein-coding genes as well as 1,519 and 1,976 pseudogenes in the WT and BR genomes, respectively (Table 1). The vast majority (15,565 and 14,727) of these genes and all the pseudogenes (1,519 and 1,976) in the WT and BR genomes were predicted based on the CDSs in the reference genomes (reference-based), while a small portion (812 and 683) of the genes were predicted based on the RNA-seq datasets in the two species (RNA-based) (Table 1). To evaluate the effects of the unbalanced RNA-seq datasets on the numbers of annotated genes in the two genomes, we repeated the annotation process by 100 times with each time using the CDSs in the 53 Aves genomes for reference-based annotations and a total of six RNA-seq datasets for RNA-based annotation. Of the six RNA-seq datasets, three were those from the BR tissues and the other three were randomly selected in each annotation from the 20 datasets of the WT tissues. As shown in Table S2, in each of the 100 times of annotations, the total number of annotated genes in WT is larger than that in BR, which is mainly due to the larger number of genes annotated by the reference-based approach in WT (15,565) than in BR (14,727). On the other hand, the two genomes have a similar average number (350 vs 355) of genes annotated by the RNA-based approach (p-value = 0.33, two-tailed t-test). Interestingly, in 73 of the 100 times of annotations, the BR genome has even more RNA-annotated genes than does the WT genome, suggesting that the assembly quality of the CDSs in BR is good enough for finding genes. However, when the union of the 100 annotations is taken, WT still has more RNA-based annotated genes (812 vs 683). Taken together, these results suggest that the difference in the numbers of genes in the two species is not mainly due to the difference in the quality of genome assemblies and/or the bias of RNA-seq datasets.

**Table 1:** Gene annotation of two *Crossoptilon* species, *C. crossoptilon* (WT) and *C. mantchuricum* (BR), using reference- and RNA-based methods.

| Breeds | Reference-based |  |  | RNA-based |  | # Total genes | # Total pseudogenes |
| --- | --- | --- | --- | --- | --- | --- | --- |
|  | # Intact | # Partial | # Pseudogenes | # NT-supported | # Novel |  |  |
| WT | 14,815 | 750 | 1,519 | 751 | 61 | 16,377 | 1,519 |
| BR | 14,518 | 209 | 1,976 | 631 | 52 | 15,410 | 1,976 |
| Common | 14,057 |  | 1,040 | 422 | 28 | 14,507 | 1,040 |

Most of the reference-based genes (14,815 and 14,518) in both species have an intact open reading frame (ORF) (intact genes), while the remaining small numbers (750 and 209) contain either premature stop codon or ORF-shift mutations, which are not supported by the corresponding short DNA reads, we thus called them partially supported genes, as the mutations might be caused by sequencing errors, particularly, in long reads. Vast majority of the intact genes (99.01%), partially supported genes (97.73%), and pseudogenes (95.13%) in WT were transcribed in at least one of the 20 tissues examined (Table S3). These percentages (96.54%, 93.78% and 90.84%) are smaller in BR (Table S4) as RNA-seq data from only three tissues are available^45^. Most (751 and 631) of the RNA-based genes in both species can be mapped to the NT database^48^ (NT-supported new genes), suggesting that they are likely true genes. The remaining small numbers (61 and 52) that cannot be mapped to the NT database are likely novel genes (Table 1).

We compared the RNA-based genes annotated in the WT and BR assemblies, and found 450 pairs with an identity > 96.8%, comprising 55.4% and 65.9% of their RNA-based genes, respectively. They are likely true orthologous genes in both species, as the same patterns of transcriptional noise are unlikely to occur in two different species. Moreover, most of the RNA-based genes in both species (Tables S3 and S4) were varyingly expressed in different tissues with RNA-seq data available (Figures 1a-1d), indicating that they might be authentic and functional. In addition, we randomly selected 17 of the 812 RNA-based genes in WT and measured their expression levels in 16 tissues using RT-qPCR. We found that 15 (88.24%) of them were transcribed in at least one of the tissues and five of which were putative novel genes (Table S5, Figure 1e), further suggesting that most of the RNA-based genes at least in WT are likely authentic, although the expression patterns of some genes are different from those seen in the RNA-seq data due probably to the different sensitivity of RT-qPCR and RNA-seq methods (Figure 1f). Moreover, we annotated 12 and eight in WT and BR, respectively, of the 274 presumed missing genes in avian species^49^ (Table S6).

**Figure 1.**
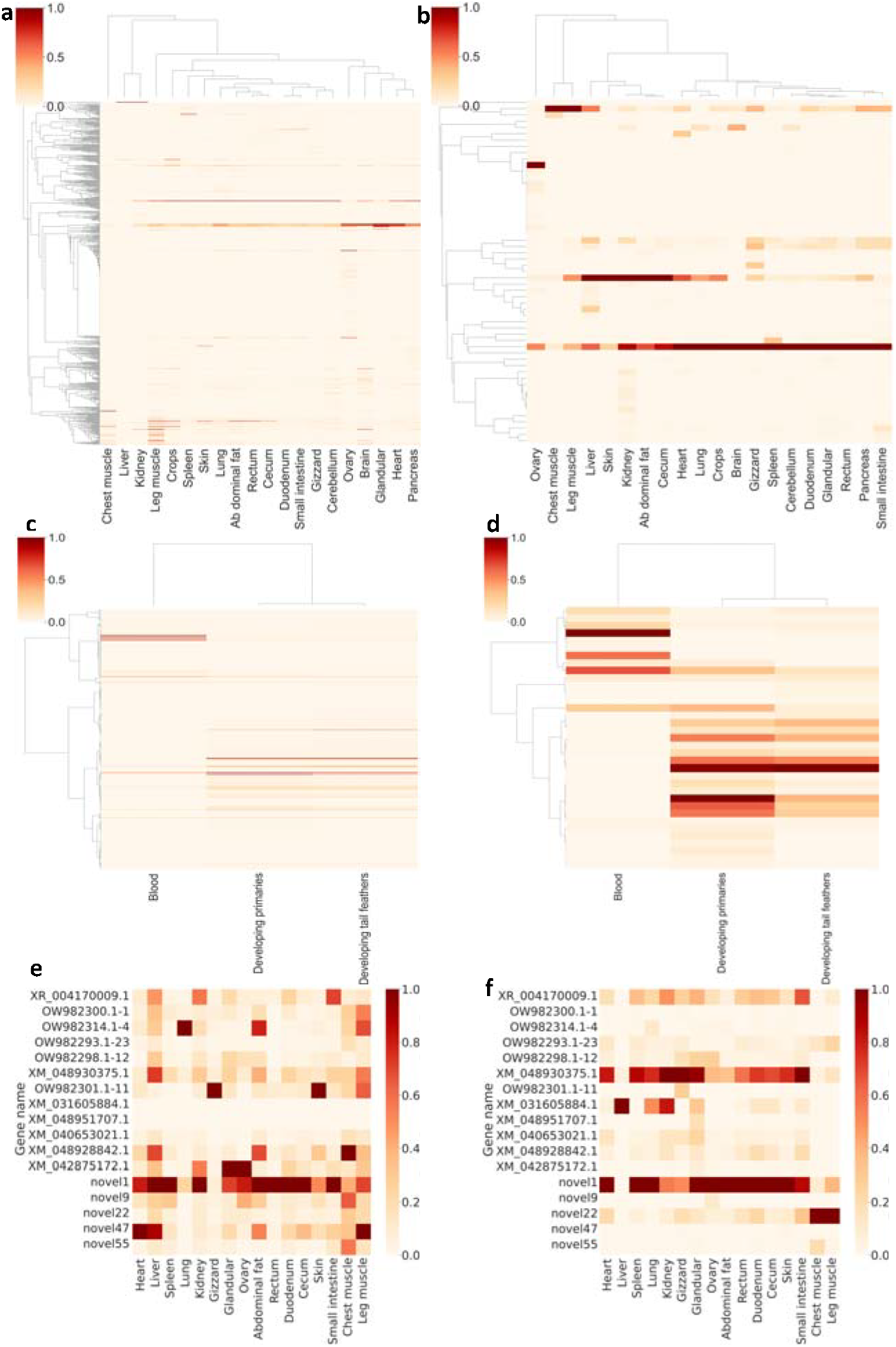
Heatmaps of expression levels of new genes in tissues of C. crossoptilon (WT) and C. mantchuricum (BR). **a.** Expression pattern of the NT-supported new genes in 20 tissues of WT. **b**. Expression pattern of the novel genes in 20 tissues of WT. **c**. Expression pattern of the NT-supported new genes in three tissues of BR. **d**. Expression pattern of the novel genes in three tissues in BR. **e**. Expression pattern of the 17 selected new genes in WT measured by RT-qPCR. **f**. Expression pattern of the 17 selected new genes quantified using RNA-seq data in WT. The expression levels of genes are scaled such that 0 represents a gene that was not expressed and 1 represents a gene that has the highest expression level in the tissue (Materials and Methods). Figures a-d and f were based on TPMs, and e was based on RT-qPCR results.

### Most pseudogenes found in the WT and BR genomes are unprocessed and unitary and thus might lose gene functions

Vast majority (for WT, 1,310 or 86.2%; for BR, 1,701 or 86.1%) of pseudogenes in both WT and BR genomes are unprocessed, i.e., they arose due to direct mutations in the parent genes. We failed to find any functional copy of these unprocessed pseudogenes; thus, they are also unitary^50^. To see whether the pseudogenization alleles in the 1,519 and 1,976 pseudogenes in the WT and BR genomes, respectively, are under selection, we mapped the short reads from 10 WT individuals and 41 BR individuals to the corresponding genomes (Materials and Methods). We found that most of the first pseudogenization alleles along the parental genes were fixed in the population of the two species (Figures 2a and 2b), suggesting that these pseudogenization events might be under positive selection and thus contribute to the phenotypic traits of the two species. Interestingly, the synonymous mutations in true genes in the two species, which are generally assumed to be selectively neutral, are largely uniformly distributed along the CDSs, except at the two ends, where the numbers of synonymous mutations decrease (Figures 2c and 2d), consistent with an earlier report in chickens^51^. The reduced synonymous mutations at the two ends of CDSs suggest that they might harbor functional elements not related to their coding functions, such as transcriptional and post-transcriptional regulatory elements^52^. In contrast, pseudogenization mutations are strongly biased to the 5’-ends and 3’-ends in both species (Figures 2c and 2d). The same phenomenon was also found in the other species such as human^51,53^ and chickens^51^. Therefore, the strongly biased pseudogenization mutations to the 5’-end and 3’-end in the two species also suggest that they are under positive selection.

**Figure 2.**
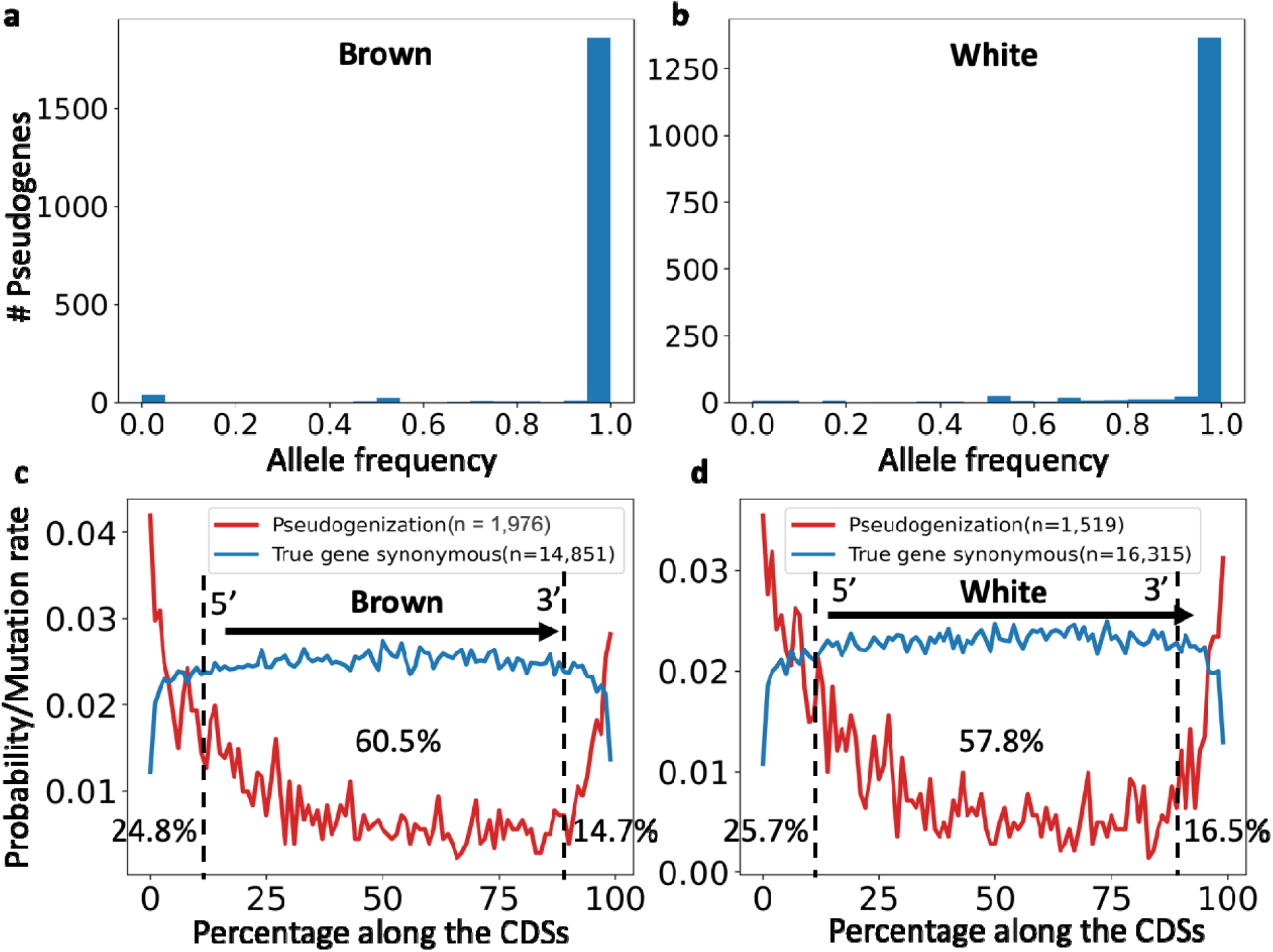
Fixation rate of pseudogenes in the WT and BR populations and distribution of the first pseudogenization alleles along the CDSs of parental genes. **a.** Number of pseudogenization alleles with indicated frequencies in the BR population. **b**. Number of the pseudogenization alleles with indicated frequencies in the WT populations. **c**. Distribution of the first pseudogenization alleles in the BR genome along the CDSs of parental genes. Notably, 24.8%, 60.5%, and 14.7% of the first pseudogenization alleles are in the first 10%, middle 80% and last 10% of the CDS regions. **d**. Distribution of the first pseudogenization alleles in the WT genome along the CDSs of parental genes. Notably, 25.7%, 57.8% and 16.5% of the first pseudogenization alleles are in the first 10%, middle 80% and last 10% of the CDS region.

Finally, of the 1,519 pseudogenes in the WT genome, 727 (47.9%) have alternative transcripts, but only 10 (1.4%) have functional alternative transcripts (Table S7). Of the 1,976 pseudogenes in BR, 1,712 have alternative transcripts, but only 17 (1.0%) have functional alternative transcripts (Table S8). These results suggest that most pseudogenes in both species do not have functional alternative transcripts, further suggesting that most of them have lost the functions of their parent genes.

### Unique genes and pseudogenes in WT and BR might be related to their adaptation to distinct ecological niches

We found that the WT and BR genomes shared 14,507 of their genes (88.6% and 94.1%) and 1,040 of their pseudogenes (68.5% and 52.6%) (Table 1). Meanwhile, the WT genome contains 1,870 unique genes and 479 unique pseudogenes, while the BR genome harbors 903 unique genes and 936 unique pseudogenes (Tables S9-S12, Figure 3a). Of the 1,870 unique genes in the WT genome, 658 (35.2%) are unique pseudogenes and 1,212 (64.8%) are missing in the BR genome. Of the 936 unique pseudogenes in BR, 658 (70.3%) are unique genes and 278 (29.7%) are missing in the WT genome (Figure 3a). Moreover, of the 903 unique genes in the BR genome, 173 (19.2%) are unique pseudogenes and 730 (80.8%) are missing in the WT genome. Of the 479 unique pseudogenes in the WT genome, 173 (36.1%) are unique genes and 306 (63.9%) are missing in the BR genome (Figure 3a). There are a total of 3,357 genes that either are unique genes or become unique pseudogenes in the WT or BR genomes. To see whether the unique genes and pseudogenes in each species are related to their unique traits and reflect the evolutionary pressure received from their niches, we assigned GO terms to the unique genes and pseudogenes in each species based on their homologs in chickens or humans. The unique genes and pseudogenes in WT are involved in 69 and 25 biological pathways (Table S13), respectively, while those in BR are involved in 27 and 59 biological pathways, respectively (Table S13). Interestingly, the unique genes and pseudogenes with GO term assignments are mainly involved in and affect, respectively, pathways related to four major physiological functions: cardiovascular functions, energy metabolism, neuronal functions and immunity (Table S13, Figure 3b). It has been shown that alterations in these functions are critical for animals to adapt to different altitudes^33^. Thus, the unique genes and loss of functions of the missing genes and pseudogenes in BR and WT might play a role in the adaption of the two species to their strikingly different altitudes. A few examples of such genes and pseudogenes are given below.

**Figure 3.**
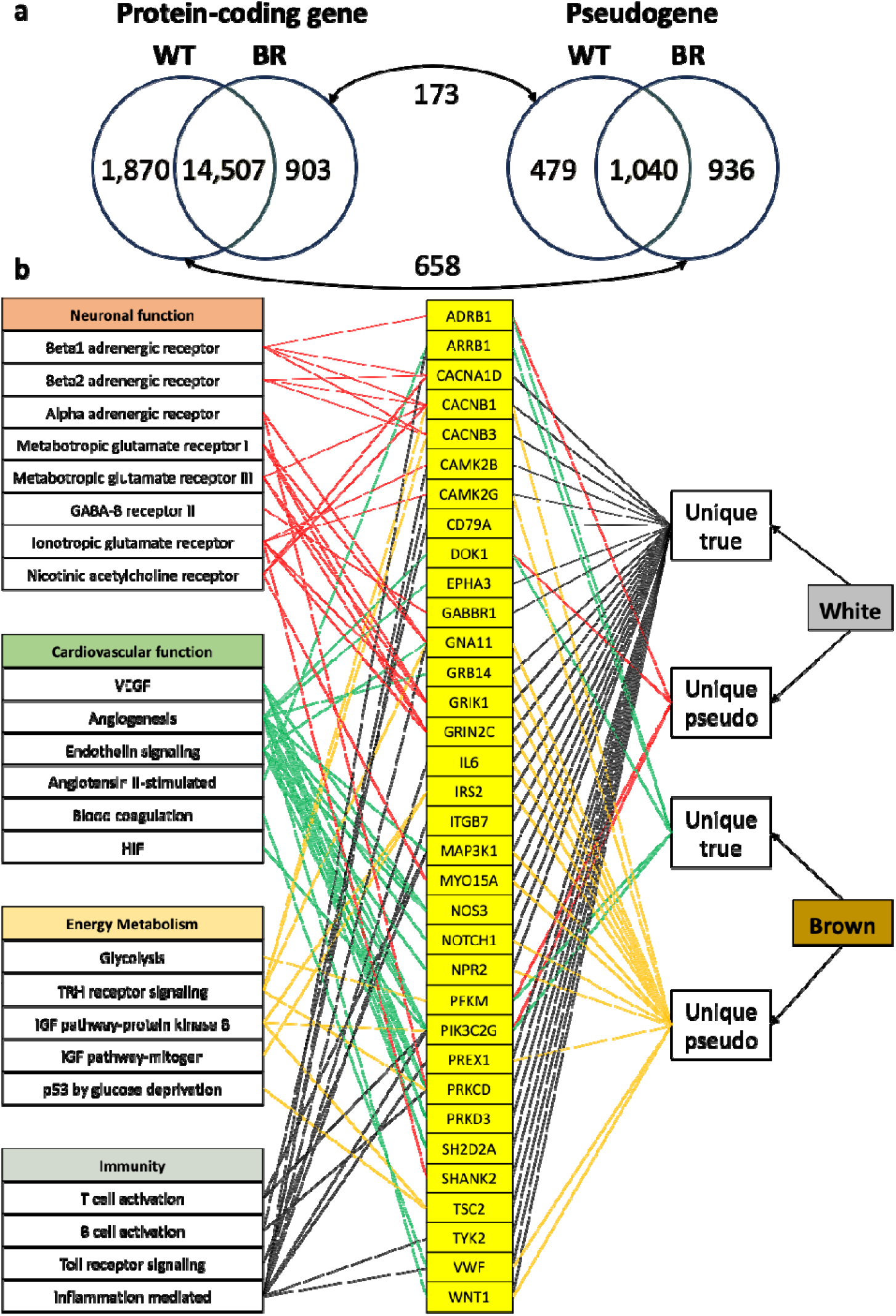
Functional analysis of the unique genes and pseudogene in the WT and BR genomes. **a.** Venn diagram showing shared and unique genes (left) and pseudogenes (right) in the two species. Of note, 658 WT genes are pseudogenized in BR and 173 BR genes are pseudogenized in WT, and there are a total of 3,357 non redundant unique genes and pseudogenes in the two genomes. **b**. The unique genes/pseudogenes in the WT and BR genomes that are involved in pathways related to four functional categories: neuronal functions, cardiovascular functions, energy metabolism, and immunity. Notably, unique genes in a species often became unique pseudogenes in the other species.

1. Cardiovascular functions: As shown in Figure 3b, unique gene *PIK3C2G* in BR, which is pseudogenized in WT, is involved in the HIF (hypoxia-inducible factor) activation pathway^54-56^. Unique genes *SH2D2A, PRKCD, PRKD3* and *NOS3* in WT, which are all missing in BR, are involved in the VEGF (vascular endothelial growth factor) signaling pathway that plays critical roles in proliferation, survival and migration of blood endothelial cells required for the angiogenesis pathway^57-59^. Unique genes *SH2D2A, MAP3K1, PRKCD, WNT1, PRKD3, NOS3, EPHA3, NOTCH1* and *GRB14* in WT, which are missing (*SH2D2A, PRKCD, PRKD3, NOS3* and *EPHA3*) or pseudogenized (*MAP3K1, WNT1, NOTCH1* and *GRB14*) in BR, are involved in the angiogenesis pathway. Unique genes *DOK1* and *PIK3C2G* in BR, which are pseudogenized in WT, are involved in the angiogenesis pathway. Unique genes *PRKCD, NOS3* and *NPR2* in WT are missing (*PRKCD* and *NOS3*) or pseudogenized (*NPR2*) in BR, are involved in the endothelin signaling pathway. Unique gene *ARRB1* in BR, which is missing in WT, is involved in the angiotensin II-stimulated signaling through G proteins and beta-arrestin pathway. Unique gene *VWF* in WT, which is pseudogenized in BR, is involved in blood coagulation pathway.
2. Energy metabolism: As shown in Figure 3b, unique gene *PFKM* in BR, which is pseudogenized in WT, is involved in glycolysis. Unique genes *PRKCD, CACNB1* and *CACNB3* in WT, which are missing (*PRKCD* and *CACNB3*) or pseudogenized (*CACNB1*) in BR, are involved in the thyrotropin-releasing hormone receptor signaling pathway. Unique genes *IRS2* and *TSC2* in WT, which are missing (*TSC2*) or pseudogenized (*IRS2*) in BR, are involved in the insulin/IGF pathway-protein kinase B signaling cascade pathway. Unique gene *TSC2* in WT, which is missing in BR, is involved in the p53 pathway by glucose deprivation.
3. Neuronal functions: As shown in Figure 3b, unique genes *CACNA1D, CACNB1* and *CACNB3* in WT, which are missing (*CACNA1D* and *CACNB3*) or pseudogenized (*CACNB1*) in BR, are members of the beta1/2 adrenergic receptor signaling pathway. Unique gene *ADRB1* in BR, which is pseudogenized in WT, is involved in the beta1/2 adrenergic receptor signaling pathway. Unique genes *GRIK1* and *GRIN2C* in WT, which are pseudogenized in BR, are involved in the metabotropic glutamate receptor group I pathway. Unique gene *CACNB1* (metabotropic glutamate receptor group III pathway) in WT is pseudogenized in BR. Unique gene *GABBR1* (GABA-B receptor II signaling pathway) in WT is missing in BR. Unique genes *CAMK2G, GRIK1, SHANK2, GRIN2C* and *CAMK2B* in WT, which are missing (*SHANK2* and *CAMK2B*) or pseudogenized (*CAMK2G, GRIK1* and *GRIN2C*) in BR, are involved in the ionotropic glutamate receptor pathway. Unique genes *CACNA1D, MYO15A* and *CACNB1* in WT, which are missing (*CACNA1D*) or pseudogenized (*MYO15A, CACNB1*) in BR, are involved in the nicotinic acetylcholine receptor signaling pathway.
4. Immunity: As shown in Figure 3b, unique gene *MAP3K1* in WT, which is pseudogenized in BR, is involved in the T cell activation pathway. Unique gene *PIK3C2G* in BR, which is pseudogenized in WT, is involved in the T cell activation pathway. Unique genes *CD79A* and *PRKCD* in WT, which are missing in BR, are involved in the B cell activation pathway. Unique genes *PREX1, CAMK2G, ITGB7, VWF, TYK2, IL6* and *CAMK2B* in WT, which are missing (*ITGB7, VWF, TYK2* and *CAMK2B*) or pseudogenized (*PREX1, CAMK2G, VWF* and *IL6*) in BR, are members of the inflammation mediated by chemokine and cytokine signaling pathway. Unique gene *ARRB1* in BR, which is missing in WT is involved in inflammation mediated by chemokine and cytokine signaling pathway.

In addition, some unique genes or unique pseudogenes in the two species are involved in pathways that might be related to the adaptation of the two species to other aspects of their distinct ecological niches. For example, unique genes *CACNA1D, PRKCD, CACNB1* and *CACNB3* in WT are missing (*CACNA1D, PRKCD* and *CACNB3*) or pseudogenized (*CACNB1*) in BR (Table S13).

These genes are involved in oxytocin receptor mediated signaling pathway that has been shown to play critical roles in social behaviors of animals^60^. Moreover, unique genes *IRS2, SMAD4, FOSB, CACNA1D, MAP3K1, PRKCD, MAP2K7, EP300, NR5A1, NPR2, MAP3K5* and *CAMK2B* in WT are missing (*SMAD4, FOSB, CACNA1D, PRKCD, MAP2K7, NR5A* and *CAMK2B*) or pseudogenized (*IRS2, MAP3K1, EP300, NPR2* and *MAP3K5*) in BR (Table S13), these genes are in involved in gonadotropin releasing hormone receptor pathway that regulate reproduction.

### The populations of the three species are strongly structured

To further understand the genetic basis of altitude adaptation, we next compared the SNPs of 10 WT individuals, 41 BR individuals and 12 BL individuals (Table 2). Figure 4a shows the geographical locations where the populations were sampled. Specifically, the three BR subpopulations were from Shaanxi province, Shanxi province, and Hebei province & Beijing metropolitan, respectively^45^; the BL population were from Gansu province^45^; and the WT population were from Yunnan province. Using our assembled WT genome^61^ as the reference, we identified SNPs in the BL, BR and WT populations (Table 2). Based on the bi-allelic SNPs (16 million) in the populations of the three species (Table 2), we performed principal component analysis (PCA), and found that individuals of the same species form distinct clusters (Figure 4b). Notably, each of the three BR subpopulations forms a distinct compact subcluster, which is consistent with the previous result^45^. We also run the admixture algorithm^62^ on the SNPs to infer their ancestral relationships. For K=2, individuals of BL and BR formed a uniformly colored cluster, while WT individuals formed the other cluster sharing small portions of ancestries with the former two species (Figure 4c), suggesting that BL and BR were derived from the same ancestor, while WT branched out earlier. This result is consistent with previous phylogenetic analysis that BL and BR are more closely related to each other^43^. For K=3, individuals of each species formed an almost uniformly colored cluster, indicating their distinct recent ancestries.

**Table 2:**
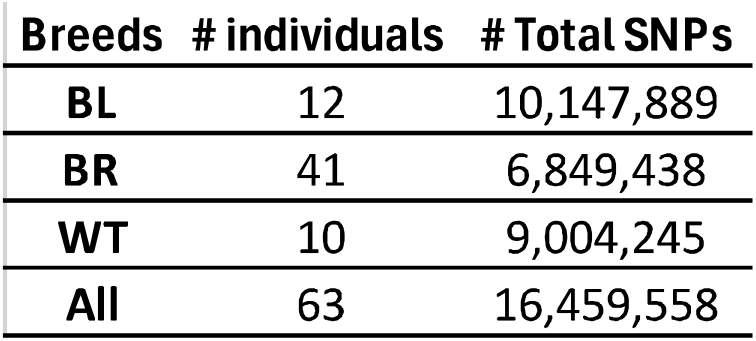
Numbers of SNPs identified in the *C. auritum* (BL), *C. mantchuricum* (BR) and *C. crossoptilon* (WT) populations.

| Breeds | # individuals | # Total SNPs |
| --- | --- | --- |
| BL | 12 | 10,147,889 |
| BR | 41 | 6,849,438 |
| WT | 10 | 9,004,245 |
| All | 63 | 16,459,558 |

**Figure 4.**
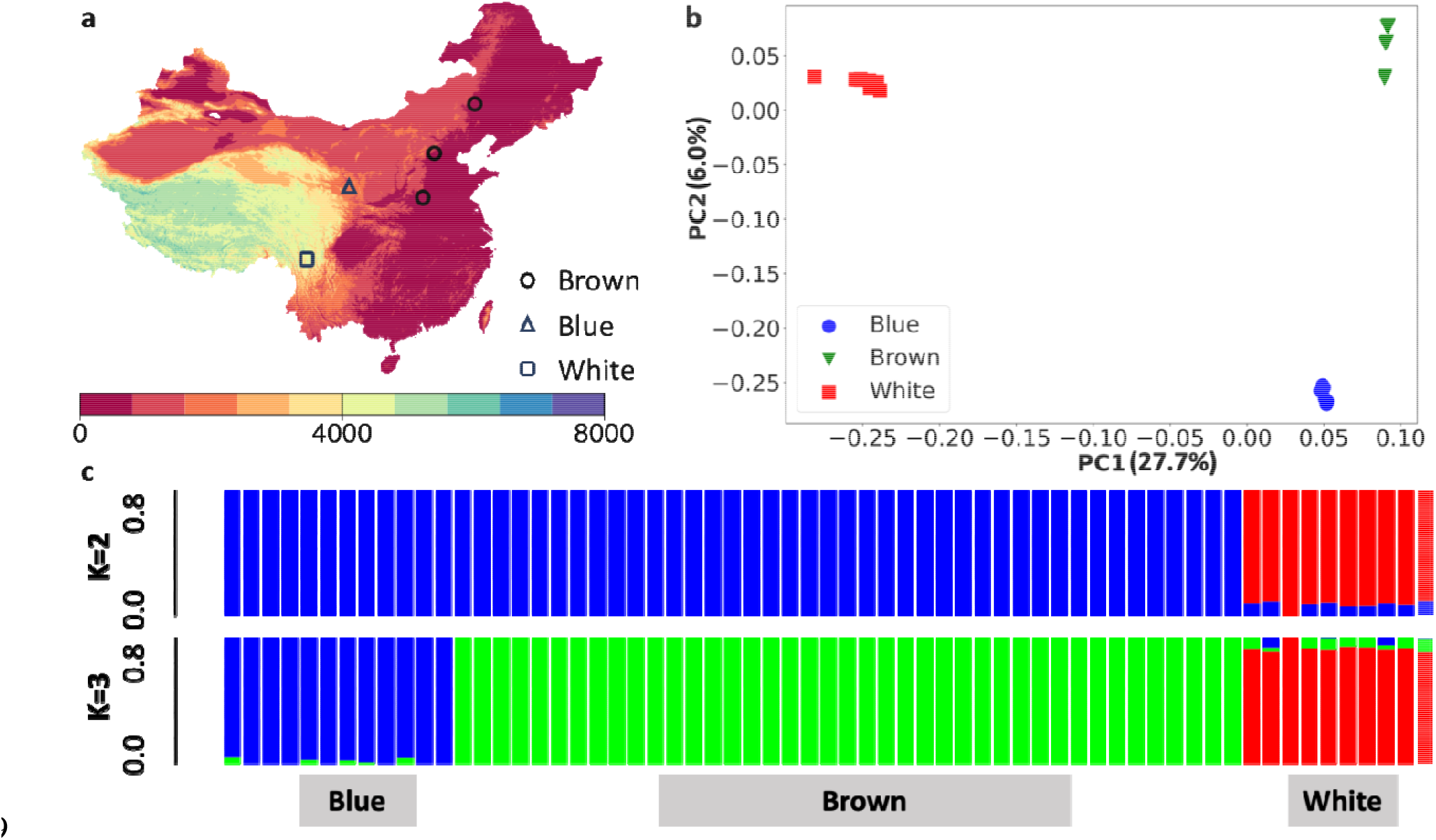
Geographic distribution and population structure of the WT, BL and BR populations. **a.** Geographic distribution of the three species used in this study. The color of the map indicates altitude level (m). **b**. Principal component analysis of the populations of the three species based on their SNP spectrum. **c**. Ancestral analysis of the individuals of the three species using ADMIXTURE for K=2 and 3.

To see possible gene introgression among the three species, we calculated the D-statistic^63^ and f3-statistic^64^. We obtained a D value of 0.57 (z-score = 36.18, and p-value= 2.3e-16) using a Daweishan chicken population as the outgroup (O) on the rooted tree (((BR,BL),WT),O), suggesting introgression between BL and WT^63^. The f3 values and associated z-scores for the three choices of (target; source 1, source 2) are shown in Table 3. The significant (z-score = −26.45) negative f3 value (−0.0586) for (target=BL; source 1 = WT, source 2 = BR) suggests introgression in BL from both WT and BR^64^. However, the positive f3 values for the other two scenarios (Table 3) could not indicate there is gene introgression in BR from WT and BL, and in WT from BR and BL^64^. Thus, although gene introgression might have occurred in BL from both BR and WT, the data does not support gene introgression in BR from WT and BL, and in WT from BR and BL. Thus, gene introgression might play a role in the evolution of BL, but this might not be the case for BR and WT.

**Table 3:** The f3-statistic and associated z-scores for the three choices of (target; source 1, source 2) among the three species.

| Source 1 | Source 2 | Target | f3 | std. err | Z-score |
| --- | --- | --- | --- | --- | --- |
| WT | BR | BL | -0.0586 | 0.0022 | -26.45 |
| WT | BL | BR | 1.2340 | 0.0769 | 16.04 |
| BR | BL | WT | 0.8392 | 0.0047 | 180.24 |

### Genes in selective sweeps of each species might be related to their distinct ecological niches

To identify genomic regions and genes that might be related to the adaptation of the three species to their distinct ecological niches, particularly, to different altitudes, we identified selective sweeps in each species by comparing its SNPs with those of the other two species using both the genetic differentiation (*F*_*ST*_) and difference in nucleotide diversity (Δ*π*) parameters^65-68^. To rigorously evaluate the statistical significance of the *F*_*ST*_ and Δ*π* values, we normalized them (*ZF*_*ST*_ and *Z*Δ*π*) using a permutation test ^69^ (Materials and Methods). Figures 5a-5c show the distribution of *ZF*_*ST*_ and *Z*Δ*π* as well as their values for each 40 kb genome windows with a 20 kb-step size for the three pairwise comparisons. We consider a window with *ZF*_*ST*_ > 2.33 (P=0.01) and *Z*Δ*π* > 1.64 (P=0.05) or *Z*Δ*π* < −1.64 (P=0.05) as a selective sweep (Figures 5a-5c). Figures 6a-6f show the Manhattan plots of Z*F*_*ST*_ and Z|Δ*π*| scores for the three pairwise comparisons. Since adjacent selective sweeps can overlap with one another, we merged the overlapping ones as a discrete selection sweep (DSS) in each species. For the BL vs BR comparison, we identified 208 and 638 selective sweeps, and 128 and 463 DSSs containing 143 and 474 genes in BL and BR, respectively (Tables 4 and S14). For the BL vs WT comparison, we identified 92 and 274 selective sweeps, and 72 and 181 DSSs containing 67 and 167 genes in BL and WT, respectively (Tables 4 and S15). For the BR vs WT comparison, we identified 306 and 359 selective sweeps, and 184 and 188 DSSs containing 196 and 167 genes in BR and WT, respectively (Tables 4 and S16).

**Table 4:**
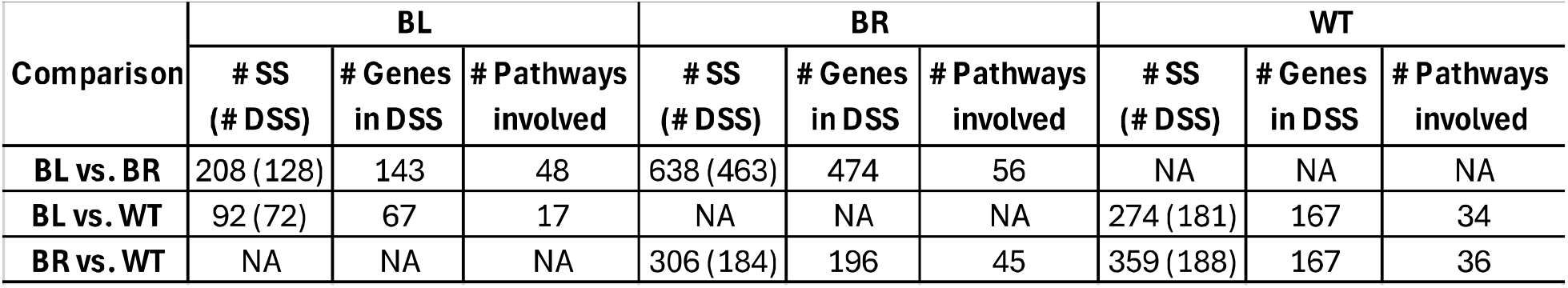
Summary of selective sweeps (SS) and discrete selective sweeps (DSS) identified in pairwise comparisons between three *Crossoptilon* species populations.

| Comparison | BL |  |  | BR |  |  | WT |  |  |
| --- | --- | --- | --- | --- | --- | --- | --- | --- | --- |
|  | # SS (# DSS) | # Genes in DSS | # Pathways involved | # SS (# DSS) | # Genes in DSS | # Pathways involved | # SS (# DSS) | # Genes in DSS | # Pathways involved |
| BL vs. BR | 208 (128) | 143 | 48 | 638 (463) | 474 | 56 | NA | NA | NA |
| BL vs. WT | 92 (72) | 67 | 17 | NA | NA | NA | 274 (181) | 167 | 34 |
| BR vs. WT | NA | NA | NA | 306 (184) | 196 | 45 | 359 (188) | 167 | 36 |

**Figure 5.**
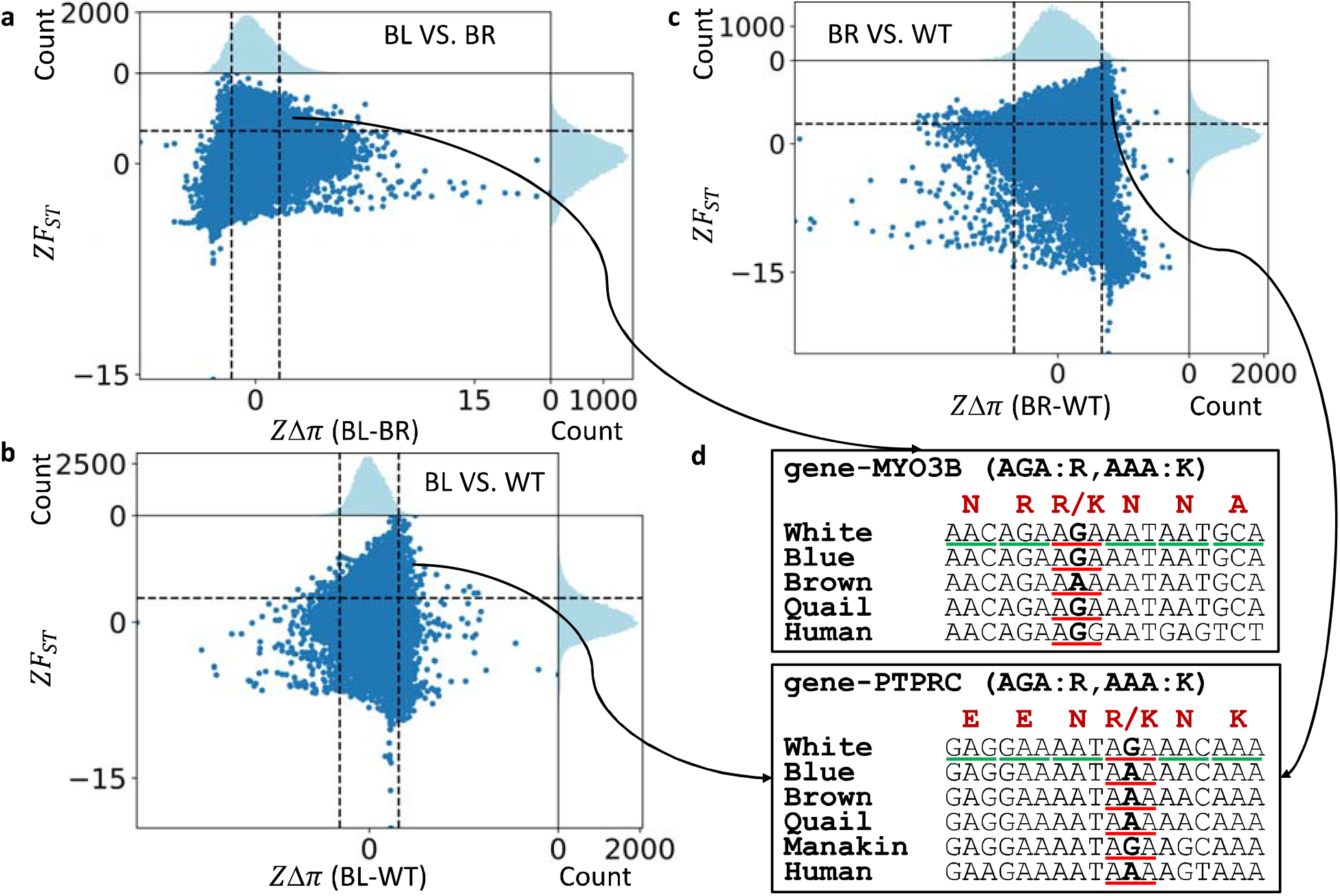
Distribution of the *ZF*_*ST*_ and *Z*Δ*π* values of each sliding window for the three pairwise comparisons. **a.** BL VS. BR comparison. **b**. BL VS. WT comparison. **c**. BR VS. WT comparison. **d**. Multiple alignments of part of CDSs of two genes in DSSs in different species. In subfigures a, b and c, the top panel represents the distribution of *Z*Δ*π* values and the bottom right one represents the distribution of *ZF*_*ST*_values. Each dot in the bottom left panel represents a sliding window with its *Z*Δ*π* and *ZF*_*ST*_ values being its coordinates in the plot. The dashed lines represent the cut-offs for the *ZF*_*ST*_ and *Z*Δ*π* values.

**Figure 6.**
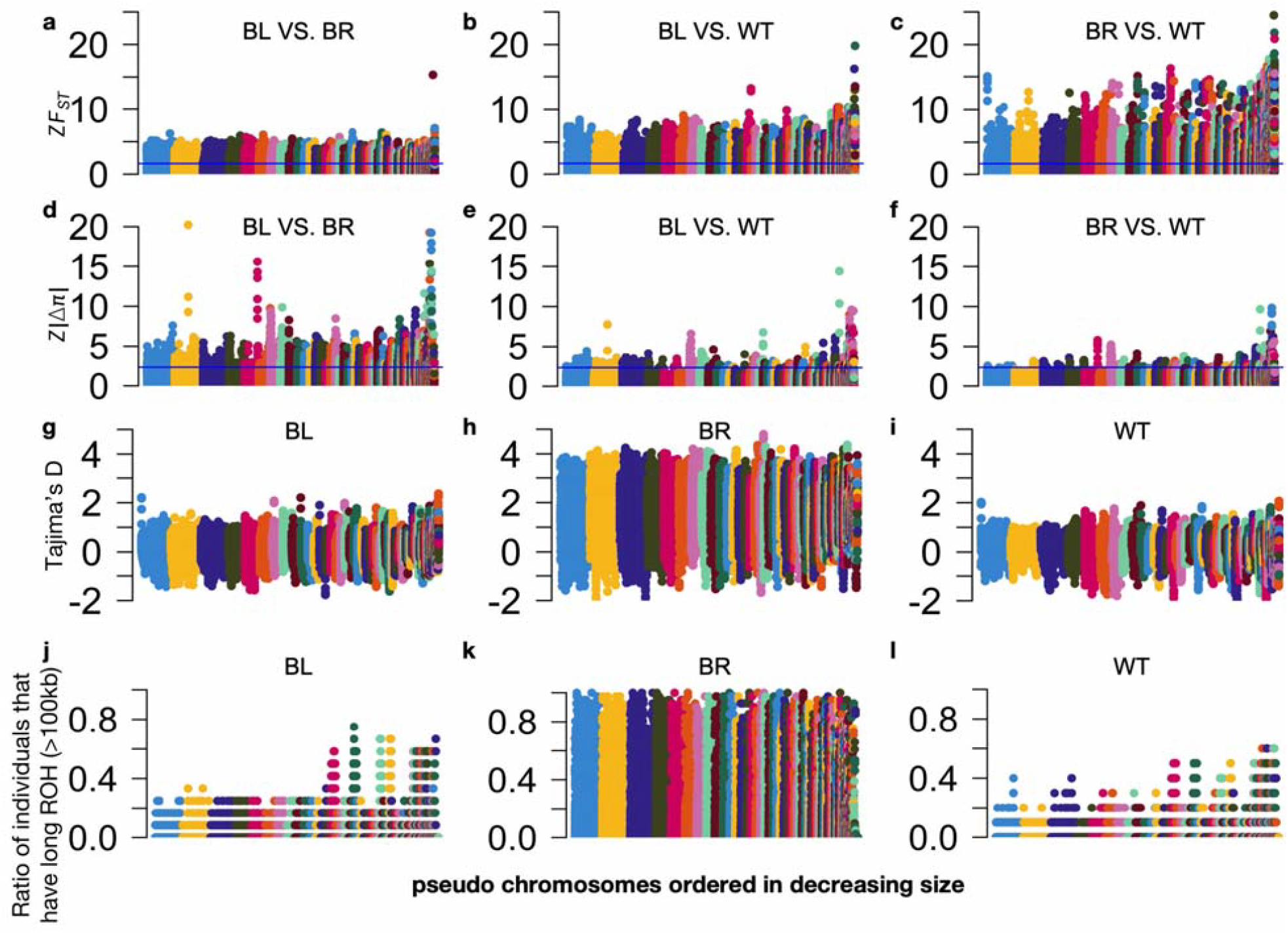
Manhattan plots for genome-wide analyses of the evolution of the three species. **a-c.** Manhattan plots of *ZF*_*ST*_ for the three pair-wise comparisons of the species. The blue line indicates the cut-off value. **d-f**. Manhattan plots of *Z*|Δ*π*| for the three pair-wise comparisons of the species. The blue line indicates the cut-off value. **g-i**. Manhattan plots of Tajima’s D values for each species. **j-l**. Manhattan plots of the ratio of individuals that have long ROH (>100kb) for each species. In the figures, each dot represents a sliding window. The horizontal axes are pseudo chromosomes ordered in decreasing size.

To further confirm our identified selective sweeps, we computed Tajima’s D values for each 40 kb window. As shown in Figures 6g-6i and Table S17, BR has the largest number (38,488) of windows with positive Tajima’s D values among the three species, suggesting that BR has experienced a large-scale population contraction, consistent with the previous report^45^. For BL and WT, most of their selective sweeps identified above in each pair-wise comparison (78.8%, 53.3%, 90% and 87.2% for BL vs BR, BL vs WT, BL vs WT and BR vs WT, respectively) have a negative Tajima’s D value (Table S17), suggesting negative or positive selections and confirming the selective sweeps identified above.

To find the duration and intensity of the selective sweep in each species, we calculated the run of homozygosity (ROH) in each selective sweep for each pair of comparison. As shown in Figures 6j-6l and Table S17, BR has the largest number of selective sweeps (66.7%-70.5%) with a high ratio of individuals that have a long ROH (>100kb), suggesting that BR has the most recent inbreeding, their selection sweeps arose recently, and the selection intensity of BR is the largest compared to the other two species. By contrast, both BL and WT have only small portions (12%-16.8%) of selective sweeps with a long ROH (Table S17). These results suggest that BR has undergone the most intensive selection recently compared to BL and WT, which is consistent with the previous report^45^.

To see whether genes in the selective sweeps are under positive or negative selections, we calculated the ratio of the number of nonsynonymous mutations over the number of synonymous mutations (dN/dS) for each gene in the DSSs for each pair of comparison. As shown in Table S17, of the genes in the identified DSSs in each pair of comparison, 76.1-90.4% are under purifying selection (dN/dS<1), while 6.6-19.4% are under positive selection (dN/dS>1). Thus, most (90.5-97.6%) of the genes in the identified DSSs are under selection.

Genes in the DSSs of each comparison might be related to adaptation to their different ecological niches. About 55% of genes in these selective sweeps identified in each comparison do not harbor fixed nonsynonymous mutations, thus, natural selection might act upon their regulatory sequences, thereby changing their expression levels. The remaining about 45% of the genes contain fixed nonsynonymous mutations that might alter the functions of encoded proteins. For example, gene *MYO3B* is located in a selective sweep in the BR population based on the BL vs BR comparison (Figure 5a). *MYO3B* in BR contains a fixed G-to-A nonsynonymous mutation in exon 7, leading to an Arg-to-Lys substitution. Since the AGA codon for Arg at this position in *MYO3B* is shared by WT, BL, quail (*Coturnix japonica*) and human (Homo sapiens) (Figure 5d), the G-to-A mutation is BR lineage specific. Thus, the Arg-to-Lys substitution in *MYO3B* in BR might be related to their unique traits. It is reported that gene *MYO3B* is required for normal cochlear hair bundle development and hearing^70^. As BR has been hunted by human for hundreds of years^48^, it is highly likely that the mutation might increase their auditory sensation, thereby helping them to escape earlier when perceiving hunters approaching. It is interesting to investigate the effects of the Arg-to-Lys mutation on the auditory functions of BR and its implication for natural selection and adaptation. Moreover, gene *PTPRC* is located in a selective sweep in the WT populations based on both the BL vs WT and the BR vs WT comparisons. *PTPRC* in WT contains a fixed A-to-G nonsynonymous mutation in exon 7, resulting in a Lys-to-Arg substitution. As the AAA codon for Lys at the position in *PTPRC* is shared by BL, BR, quail and human (Figure 5d), the mutation is WT lineage specific. Interestingly, we found that the same mutation existed in the *PTPRC* genes of Lance-tailed manakin (*Chiroxiphia lanceolata*) and White-ruffed manakin (*Corapipo altera*) (Figure 5d), both are altitudinal migrants living in subtropical Andes highlands^71^, due probably to convergent evolution. It has been reported that *PTPRC* is related to the T cell activation^72^. Therefore, it is highly likely that the nonsynonymous mutation in the *PTPRC* gene in WT and manakins might help them adapt to high altitudes where there are fewer pathogens, and thus, the pressure on the immune system in these species might be relaxed.

### Genes under selection tend to be involved in the same altitude adaptation-related pathways in the three species

There are a total of 1,106 genes that are in a selective sweep predicted based on at least one of the three comparisons. They share only 105 genes with the 3,357 unique genes and pseudogenes in the WT and BR genomes (Figure 7a). Intriguingly, the genes with GO term assignments in the selective sweeps in each species tend to be involved in the same pathways in the four functional categories (Tables S14-S16) as are the unique genes and unique pseudogenes with GO term assignments in the WT and BR genomes (Table S13), although the two sets only share four genes (Figure 7b). For example, selection on cardiovascular function-related genes *MAPK1* and *PRKAR2B* in BL might be related to its adaptation to intermediate altitudes, while selection on neuronal function related gene *PRKAR2B* might be beneficial for BL to survive in its ecological niches (Figure 7b). Moreover, selection on cardiovascular (*PLA2G4A*), immunity (*ITPR2*), energy metabolism (*PRKCD*) and neuronal function (*SLC17A6*) related genes might be beneficial for BR to adapt to low altitude (Figure 7b). Furthermore, selection on the hypoxia response (*PPARG*) related genes might help WT to adapt to the high-altitude niches, while selection on immunity (*PPP3CB, MAPK8* and *PTPRC*) and neuronal function (*GRIN3A*) related genes might be beneficial for WT to adapt to their unique ecological niches (Figure 7b). More examples of genes in the selective sweeps, which are involved in altitude-adaptation related pathways, are given as follows.

**Figure 7.**
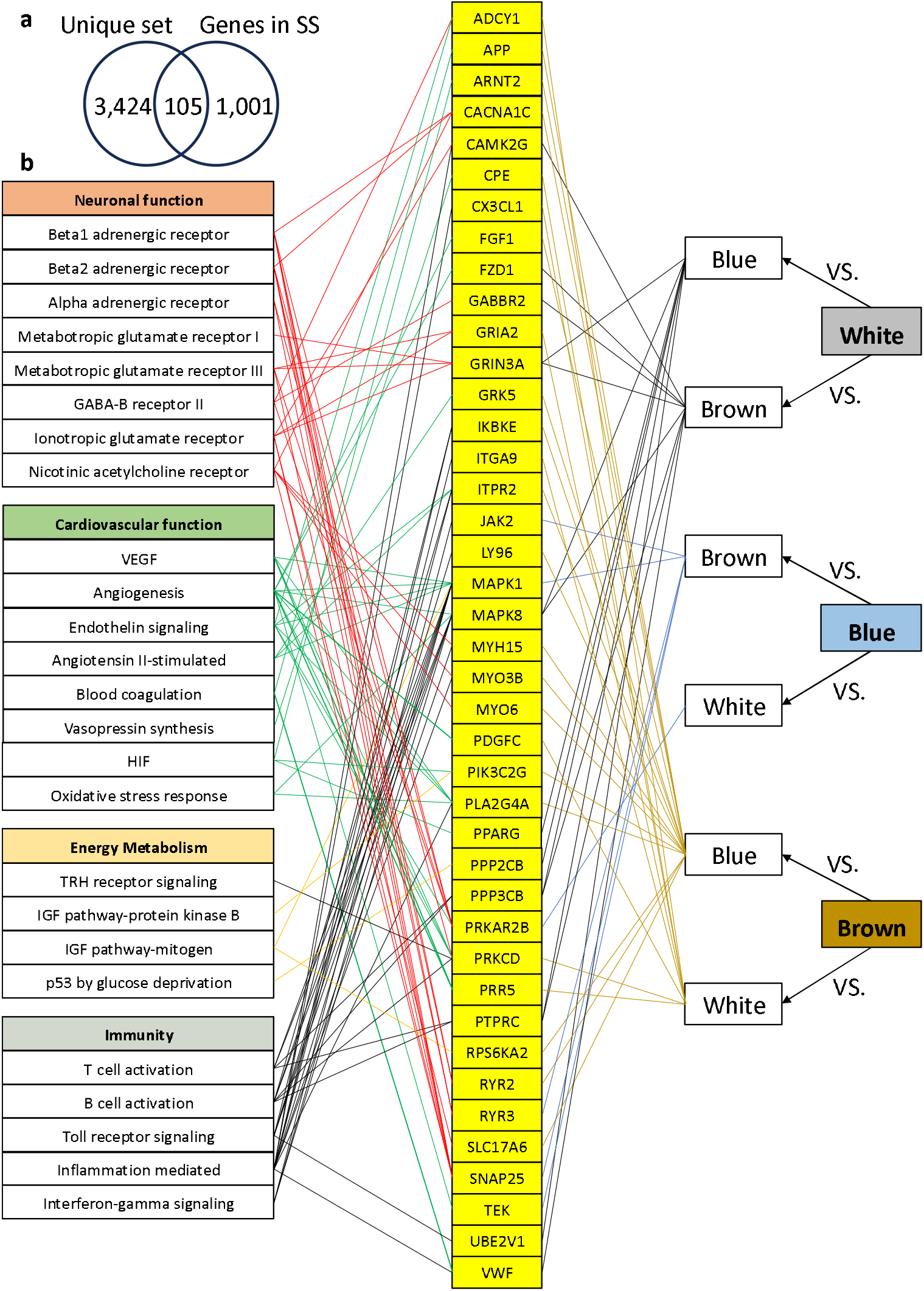
Comparison of unique genes and pseudogenes in the WT and BR genomes and genes in selective sweeps in the three species. **a.** Venn diagram of the set of unique genes and pseudogenes in the WT and BR genomes (unique set) and the set of genes in selective sweeps in the three species (genes in SS). **b**. The genes with GO term assignment in selection sweeps in each species that are involved in the same pathways of the same four functional categories as are the unique genes and unique pseudogenes with GO term assignment in the WT and BR genome, although the two sets only share four genes, i.e., *CAMK2G* and *VWF* are pseudogenized in BR, *PRKCD* is missing in BR, and *PIK3C2G* is pseudogenized in WT (Figure 3b).

For BL, genes in selective sweeps identified in the BL vs BR and BL vs WT comparisons are involved in the same number of 22 GO pathways (Table 4). Compared to BR, six genes in the selection sweeps in BL have GO pathway assignments (Table S14). Among these genes, *MAPK1* is involved in cardiovascular functions such as angiogenesis pathway, VEGF signaling pathway^57-59^, endothelin signaling pathway^73-75^, and angiotensin II-stimulated signaling through G proteins and beta-arrestin pathway (Figure 7b). Compared to WT, 76 genes in the selection sweeps in BL have GO pathway assignments (Table S15). Among these genes, *PRKAR2B* is involved in neuronal functions such as beta1/2 adrenergic receptor signaling pathway, metabotropic glutamate receptor group III pathway, GABA-B receptor II signaling pathway and endothelin signaling (Figure 7b).

For BR, genes in the selective sweeps identified in the BL vs. BR and BR vs WT comparisons are involved in 27 and 32 GO pathways, respectively (Table 4). Compared to BL, 175 genes in the selection sweeps in BR have GO pathway assignments (Table S14). Among these genes, some are involved in cardiovascular functions, such as *PLA2G4A* (VEGF pathway and angiogenesis pathway), *PLA2G4A* and *ITPR2* (endothelin signaling pathway), *ITPR2* (angiotensin II-stimulated signaling through G proteins & beta-arrestin pathway), *PLA2G4A* (oxidative stress response pathway); some are involved in immunity such as *PLA2G4A* and *ITPR2* (B cell activation pathway); some are involved in neuronal functions, such as *RYR2* (beta1/2 adrenergic receptor signaling pathway), *SLC17A6* (ionotropic glutamate receptor pathway) (Figure 7b). Compared to WT, 78 genes in the selection sweeps in BR have GO pathway assignment (Table S16). Some of these genes are involved in cardiovascular functions, such as *PRKCD* (VEGF pathway, angiogenesis pathway, endothelin signaling pathway) that is missing in the selection sweep in BR. *PRKCD* is also involved in energy metabolism (thyrotropin-releasing hormone receptor signaling pathway), immunity (B-cell activation pathway) and neuronal functions (alpha adrenergic receptor signaling pathway) (Figure 7b). Moreover, *PRKAR2B* is also under selection in BR, which is involved in neuronal functions (Figure 7b).

For WT, genes in the selective sweeps identified in the BL vs WT and BR vs WT comparisons are involved in 62 and 48 GO pathways, respectively (Table 4). Compared to BL, 341 genes in the selection sweeps in WT have GO pathway assignments (Table S15). Of these genes, some are involved in cardiovascular functions such as *ARNT2* and *PPARG* (hypoxia induced factor pathway^54-56,76^), *FZD1* and *MAPK8* (angiogenesis pathway) and *MAPK8* (oxidative stress response pathway); some are involved in energy metabolism pathway, such as *PPP2CB* (p53 pathway by glucose deprivation); some are involved in neuronal functions, such as *GRIN3A* (metabotropic glutamate receptor group I pathway, metabotropic glutamate receptor group III pathway and ionotropic glutamate receptor pathway) and *GABBR2* (GABA-B receptor II signaling pathway); and some are involved in immunity, such as P*PP3CB, MAPK8* and *PTPRC* (T/B cell activation pathway), *MAPK8* and *UBE2V1* (toll receptor signaling pathway and interferon-gamma signaling pathway) (Figure 7b).

Moreover, compared to BR, 254 genes in the selection sweeps in WT have GO pathway assignments (Table S16). Of the these genes, some are involved in cardiovascular functions, such as *MAPK8* (oxidative stress response pathway^77-81^), *TEK, FZD1* and *MAPK8* (angiogenesis pathway), and *VWF* (blood coagulation pathway) that is missing in BR (Figure 3b); Some are involved in immunity, such as *MAPK8* (interferon-gamma signaling pathway and toll receptor signaling pathway), *PPP3CB, MAPK8* and *PTPRC* (T/B cell activation pathway and toll receptor signaling pathway and interferon-gamma signaling pathway^82-84^), *CAMK2G* (pseudogenized in BR) and *VWF* (inflammation mediated by chemokine and cytokine signaling pathway^85,86^); some are involved in neuronal functions, such as *GRIN3A* (metabotropic glutamate receptor group I pathway, metabotropic glutamate receptor group III pathway), *GABBR2* (GABA-B_receptor_II_signaling pathway), *CAMK2G* and *GRIN3A* (ionotropic glutamate receptor pathway) (Figure 7b).

## Discussion

To better understand the genetic bases of adaptation of closely related eared pheasant species to highly varying altitudes, we combined our own published data in WT and BL^46^ with those in BR and BL published by Wang et al.^45^. We first annotated the assembled WT^46^ and BR^45^ genomes, and compared their gene compositions. We identified similar numbers of protein-coding genes in the WT (16,447) and BR (15,410) genomes to those annotated in other avian species^87^, but surprisingly large numbers of pseudogenes in them (1,519 in WT and 1,976 in BR), not previously reported in other avian genomes, to our best knowledge. We provided evidence that most pseudogenization mutations are under positive selection and the resulting pseudogenes might have lost gene functions.

We found that many unique genes in one species (WT or BR) are often either missing or pseudogenized in the other species (BR or WT). Interestingly, these unique genes and genes with loss-of-function mutations (pseudogenization or missing) are mainly involved in and affect, respectively, pathways related to four functional categories: cardiovascular, neuronal, energy metabolic, and immune functions. This finding is quite interesting but not very surprising, since cardiovascular, neuronal, and metabolic functions are highly sensitive to blood oxygen levels which is directly affected by oxygen pressures at varying altitudes. Moreover, varying pathogens and ultraviolet radiations at varying altitudes might require different immune responses to infections and DNA damages. Physiological systems often contain both positive and negative regulatory pathways to achieve a homeostasis of the functions. Thus, both the presence of a gene involved in the positive pathway and the loss of a gene involved in the negative pathway can enhance the relevant function. Conversely, both the presence of a gene involved in the negative pathway and the loss of gene involved in positive pathway can suppress the relevant functions. More specifically, the presence or the loss of genes in WT might lead to changes in these four functional categories that are beneficial to its adaptation to high altitude, as have been reported for other high-altitude adapted species and populations of humans and animals^4,7,11,15,18,19,22,23,31^. And the loss or the presence of the same genes in BR might lead to opposite changes of these four functional categories that are beneficial to its adaptation to low altitude. For example, we found that gene *PIK3C2G* in BR is pseudogenized in WT. This gene is involved in many pathways, including the HIF pathway, endothelin signaling pathway, angiogenesis pathway, VEGF signaling pathway, T cell activation pathway and inflammation mediated by chemokine and cytokine signaling pathway (Figure 3b). It is likely that functional *PIK3C2G* in BR plays a role in BR’s possibly low demands for these cardiovascular and immune functions to adapt to low altitudes, while the loss-of-function of *PIK3C2G* in WT might better meet WT’s possibly high demands for these functions to adapt to high altitudes. Moreover, gene *CACNB1* in WT is pseudogenized in BR. This gene is involved in the beta1/2 adrenergic receptor signaling pathway, metabotropic glutamate receptor group III pathway, nicotinic acetylcholine receptor signaling pathway and thyrotropin-releasing hormone receptor signaling pathway (Figure 3b). It is likely that functional *CACNB1* in WT might play a role in WT’s possibly high demands for these neuronal and energy metabolic functions to adapt to high altitudes or other aspects of its unique niches, while the loss-of-function of *CACNB1* in BR might better meet BR’s possibly low demands for these functions to adapt to low altitudes or other aspects of its unique niches.

Next, we compared the SNPs in the WT, BL and BR populations. Interestingly, we found that genes in the selective sweeps in each species are mainly involved in the same GO pathways of the four functional categories as are the unique genes and pseudogenes in the WT and BR genomes, even though the two sets of genes have little overlaps (Figure 7a). Most genes in the selective sweeps do not have amino acid-altering mutations, although some may have missense mutations such as *MYO3B* in BR and *PTPRC* in WT (Figure 5d), become pseudogenes such as *CACNB2*, or be missing such as *PRKCD* in BR. These results suggest that natural selection might act on their cis-regulatory sequences, thereby altering their expression levels in relevant tissues. It appears that natural selection on these genes might be related to the adaptation of the species to different niches, particularly different altitudes. For example, in the BL vs BR comparison, gene *MAPK1* in BL is included in a selective sweep. This gene is involved in the VEGF signaling pathway, angiogenesis pathway, endothelin signaling pathway, angiotensin II-stimulated signaling through G proteins and beta-arrestin pathway, insulin/IGF pathway-mitogen activated protein kinase kinase/MAP kinase cascade pathway, and many pathways of the immunity (Figure 7b). It is likely that selection on *MAPK1* might be related to the adaptation of BL to intermediate altitude. In the WT vs BL comparison, gene *PPARG* is included in a selective sweep, and the gene is involved in the HIF pathway^76^ (Figure 7b), thus, selection on the gene might be related to the adaptation of WT to low oxygen pressure at high altitudes.

It is not surprising that the unique genes/pseudogenes in WT and BR and genes in the selective sweeps of BL, BR and WT converge on the same sets of pathways involved in cardiovascular, energy metabolic, neuronal and immune functions. On the one hand, high altitude niches with low oxygen pressure, low temperature and less availability of food would pose a direct pressure on WT’s cardiovascular system and energy metabolism functions, while such pressure would be somewhat relaxed on BL and even more relaxed on BR. On the other hand, other altitude related ecological factors might exert different pressures on the three species for their neuronal functions for food foray and escaping for predators, and for immune functions for different pathogens at varying altitudes. Indeed, it has been shown that high altitude niches can have effects on the neuronal functions such as sensory perception in domestic yaks (*Bos grunniens*)^16^, and olfaction in Tibetan pig^7^, wild boars^7^ and ground tit (*Parus humilis*)^15^. In this study, we found that a nonsynonymous mutation in *MYO3B* in the BR population might alter their auditory sensation. It has also been shown that high altitude niches can have effects on immune functions in ground tit^15^. In this study, we found that a nonsynonymous mutation in *PTPRC* in the WT population and Manakins might alter their T-cell functions. Thus, it appears that the three eared pheasant species might adapt to highly varying altitudes or other aspects of their distinct ecological niches by loss-of-function mutations and fine-tuning genes involved in the same set of pathways to meet their varying demands for the four major functions. In other words, genes in this set of pathways might play roles in the adaptation of the three pheasant species to different niches, particularly, different altitudes by complete deletion (gene missing), pseudogenization, alteration of CDSs, or alteration of cis-regulatory sequences.

Finally, to our best knowledge, we provided thus far the most comprehensive annotation of genes and pseudogenes in the WT and BR genomes. We also identified numerous selection sweeps in the WT, BR, and BL genomes. These results can be valuable resources for researchers to better understand the biology of these uniquely adapted species and for policy makers to design strategies for the preservation of these endangered species.

## Materials and Methods

### Datasets

We downloaded our previously published data^46^, including the assembled genome of the WT individual from GenBank with accession number GCA_036346035.1, re-sequencing data of 10 WT individuals from NCBI SRA with accession number SRP433016, re-sequencing data of a BL individual from NCBI SRA with accession number SRP471294, and RNA-seq data from 20 tissues (chest muscle, leg muscle, liver, kidney, lung, spleen, heart, cerebellum, brain cortex, ovary, glandular, pancreas, abdominal fat, skin from chest, crops, rectum, cecum, duodenum, small intestine and gizzard) of the WT individual from NCBI SRA with accession number SRP432961. We downloaded the published datasets^45^, including the assembled genome of the BR individual from China National Genomics Data Center (CNGDC) with accession number GWHAOPW00000000, re-sequencing data of 41 BR individuals from CNGDC with project number PRJCA003284, RNA-seq data from three BR tissues (blood, developing primaries, and developing tail feathers) from CNGDC with project number PRJCA003284, and re-sequencing data of 12 BL individuals from CNGDC with project number PRJCA003284. We downloaded re-sequencing data of 24 individuals of an indigenous chicken (Daweishan chicken) (Gallus gallus) breed^67^ from the NCBI SRA database with project number PRJNA893352.

### Ethics approval

All the experimental procedures were approved by the Animal Care and Use Committee of the Yunnan Agricultural University (approval ID: YAU202103047). The care and use of animals fully complied with local animal welfare laws, guidelines, and policies.

### Real-time quantitative PCR (RT-qPCR) analysis

Three female adult WT individuals were collected from Diqing Tibet Autonomous Prefecture, Yannan Province, China. The birds were killed by electric shock to unconsciousness followed by neck artery bleeding. One to two grams of 20 tissues were aseptically collected from each individual bird in a centrifuge tube within 20 mins after sacrifice and immediately frozen in liquid nitrogen, then stored at −80℃ until use. The collected tissues included chest muscle, leg muscle, liver, kidney, lung, spleen, heart, cerebellum, brain cortex, ovary, glandular, pancreas, abdominal fat, skin from chest, crops, rectum, cecum, duodenum, small intestine and gizzard. Total RNA from each tissue sample was extracted using TaKaRa MiniBEST Universal RNA Extraction Kit (TaKaRa Biotechnology Co., Ltd., Dalian, P. R. China) according to the vendor’s instructions. cDNA was synthetized from the total RNA by using a PrimeScript RT reagent Kit (TaKaRa Biotechnology Co., Ltd., Dalian, P. R. China) following the vendor’s instructions as previously described^88^. RT-qPCR was performed using the Bio-Rad CFX96 real-time PCR platform (Bio-Rad Laboratories, Inc.) and SYBR Green master mix (iQTM SYBRGreen® Supermix, Dalian TaKaRa Biotechnology Co. Ltd.). We randomly selected 17 putative new genes for validation. The primers of the 17 putative new genes and their annealing temperatures are listed in Supplementary Note. The β-actin gene was used as a reference. Primers were commercially synthesized (Shanghai Shenggong Biochemistry Company, P.R.C.). Each PCR reaction was performed in 25 μl volumes containing 12.5 μl of iQ™ SYBR Green Supermix, 0.5 μl (10 mM) of each primer, and 1 μl of cDNA. Detection of products was performed with the following cycle profile: one cycle of 95 °C for 2 min, and 40 cycles of 95 °C for 15 s, the annealing temperature (Supplementary Note) for 30 s, and 72 °C for 30 s, followed by a final cycle of 72 °C for 10 min. The 2^−ΔCt^ method was used to analyze mRNA abundance. All tissues were analyzed with three biological replicates and each biological replicate with five technical replicates.

### Protein-coding gene annotation

To annotate the protein-coding genes and pseudogenes in the WT and BR genomes, we used a combination of reference-based and RNA-based method as previously described^89^. Briefly, for the reference-based method, we used the CDSs of 53 well-represented avian genomes in NCBI (Table S1) as the templates. We mapped all the CDSs isoforms of genes in the 53 genomes to the assembly using Splign (2.0.0)^90^ using default settings. For each template gene whose CDSs could be mapped to the assembly, we concatenated all the mapped parts on the assembly. If the CDSs of the template gene could be mapped to multiple loci on the assemblies, we chose the locus with the highest mapping identity and checked whether the resulting concatenated sequence formed an intact ORF (it had a length of an integer multiple of three and contained no premature stop codon). If resulting concatenated sequence formed an intact ORF, we called the locus an intact gene. If resulting concatenated sequence did not form an intact ORF, i.e. its length was not an integer multiple of three and/or it contained a premature stop codon, we mapped the Illumina DNA short reads from the same individual to the locus in the assembly allowing no mismatch and gaps using bowtie (v2.4.1)^91^ with default settings. If each nucleotide position in the locus can be mapped by at least 10 short reads, we considered the locus as a pseudogene; otherwise, we called it a partially supported gene since the pseudogenization mutation might be caused by the sequencing errors in the assembly that cannot be corrected by the polishing process.

For the RNA-based method, we first mapped all the RNA-seq reads from the 20 tissues of WT (Table S3) and the three tissues of BR (Table S4) to the rRNA database SILVA_138^92^ using bowtie (v2.4.1)^91^ with default settings, and filtered out the mapped reads. We then mapped the unaligned reads to each genome using STAR (2.7.0c)^93^ with default settings. As the *Crossoptilon* species are closely related^35,36^, here, we used a total of 23 RNA-seq datasets from both species in annotating each genome to fully utilize the available data. Particularly, the datasets of BR were collected from tissues (blood, developing primaries, and developing tail feathers) that were not collected from WT, thus, they were complementary to facilitate gene finding. Based on the mapped reads to each genome, we assembled transcripts using genome-guided method in Trinity (2.8.5)^94^ with default settings. Then we mapped the assembled transcripts to their corresponding genome using Splign (2.0.0)^90^ with default settings, and defined their gene structures, and removed those that partially overlap the genes and pseudogenes predicted by the reference-based method or the non-coding RNA genes (see below). For each unmapped assembled transcripts, we found the longest ORF in it, and called it a protein-coding gene if it was at least 300 bp long.

For completeness, we also annotated tRNA, miRNA, rRNA, snoRNA, telomerase RNA and SRP RNA genes in each genome using infernal (1.1.2)^95^ with the Rfam (v.14) database^96^ as the reference using default settings.

### Single nucleotide variants calling

To call the variants in the populations of BL, BR, WT and Daweishan chicken, we first mapped the short reads of each individual to the WT genome using BWA (0.7.17)^97^ and SAMtools (1.9)^98^ using default settings, and then called the variants for each individual using GATK-HaplotypeCaller (4.1.6)^99^ using default settings and merged the variants of each individual from the same species using GATK-CombineGVCFs (4.1.6)^99^ using default settings. We removed the SNPs with Quality by depth (QD) < 2, Fisher strand (FS) > 60, Root mean square mapping quality (MQ) < 40, Strand odd ratio (SOR) > 3, Rank Sum Test for mapping qualities (MQRankSum) < −12.5 and Rank Sum Test for site position within reads (ReadPosRankSum) < −8 and indels with QD < 2, FS > 200, SOR > 10, Likelihood-based test for the consanguinity among samples (InbreedingCoeff) < −0.8 and ReadPosRankSum < −20.

To calculate the fixation rate of the pseudogenes in the population of each species, we called the SNPs in the pseudogenes of each species using the DNA short reads from the species’ population (n=10 for WT and n=41 for BR) using the same method mentioned above. The annotated pseudogenes of each species were used as the reference for the SNP calling.

### PCA, ancestry, and introgression analyses

The called biallelic SNPs in each species located in strong linkage disequilibrium blocks and those with a MAF<0.05 were filtered. Principal component analysis (PCA) was performed on the remaining SNPs using PLINK (1.90)^100^ with default settings. The same SNPs were used to infer the ancestral relationships of the three species using ADMIXTURE (1.3.0)^62^ with default settings. The results for the number of ancestral groups K = 2 and 3 were tested. The same SNPs were used to infer possible gene introgression among the three species by computing the D-statistic using Dsuite^63^ with default settings and the f3-statistic using ADMIXTOOLS^64^ with default settings.

### Calculation of allele frequencies of pseudogenes

We computed allele frequencies of the first pseudogenization mutation of each pseudogene in each species using GATK (4.1.6)^99^ based on called SNPs and indels.

### Selective sweeps detection

Selective sweeps in a species relative to another were detected using two methods including genetic differentiation (*F*_*ST*_)^101^ and difference in nucleotide diversity (Δ*π*) ^68^. These two methods were widely used to detect selective sweeps by many studies^65-68^. We estimated *F*_*ST*_ for each pair of comparison using VCFtools (0.1.16)^102^ with a sliding window of 40 kb and a step size of 20 kb with default settings. We estimated *π* for each species using VCFtools (0.1.16)^102^ with a sliding window of 40 kb and a step size of 20 kb with default settings, and calculated the difference in nucleotide diversity (Δ*π*) in each window for each pair of comparison. To evaluate the statistical significance of the F_ST_ and π values for a comparison, we generated a Null model by shuffling the allele frequency data for 100 times while keeping the SNP positions fixed^69^. We then computed F_ST_ and Δπ for the permuted windows as well as their means 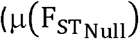 and μ(Δπ_Null_)) and standard deviations 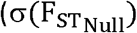 and σ(Δπ_Null_)). We computed the Z value for each F_ST_ and Δπ value for the comparison by using the following formulas:

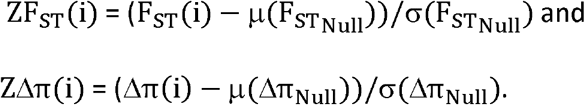

We considered a window with *ZF*_*ST*_ > α and *Z*Δ*π* > β or *Z*Δ*π* < -β (α>0, β>0) as a selective sweep. For the selective sweeps in each pair of comparison, we used the value of *Z*Δ*π* to help identify the selective sweeps of each species. Windows with *Z*Δ*π* scores < -−β represent selective sweeps of the minuend species, and windows with *Z*Δ*π* scores > β represent selective sweeps of the subtrahend species. We set α = 2.33 and β = 1.64, corresponding to a one-tailed and a two-tailed z-test, respectively.

### Runs of homozygosity (ROH) analysis

ROH analysis in each species was done using BCFtools (1.10)^103^ with the default settings based on the SNPs called in each individual bird of the species.

### Tajima’s D value calculations

Tajima’s D in each species was calculated using ANGSD (0.930)^104^ with default settings based on the SNPs called in each individual bird of the species.

### RNA-seq data analyses

RNA-seq reads of each tissue were mapped to the WT or BR genome using STAR (2.7.0c)^93^ with default settings. The expression level of each gene or pseudogene *g*_*i*_ in the tissue was computed as TPM (transcript per million), defined as 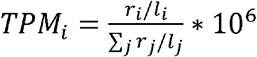, where *r*_*i*_ denotes reads mapped to *g*_*i*_, *l*_*i*_ is the length of *g*_*i*_, and ∑_*j*_ *r*_*j*_ /*l*_*j*_ corresponds to the sum of mapped reads to each *g*_*j*_ normalized by its length.

### Hierarchical clustering of gene expression levels

The expression level of each gene *g*_*i*_ in a tissue was rescaled as 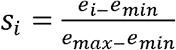,where *e*_*i*_ denotes the expression level of *g*_*i*_, and *e*_*min*_ and *e*_*max*_ the minimal and maximal expression levels of all genes in the tissue, respectively. The clustermap program from seaborn in Python was used to generate the heatmaps in Figure 1.

## Data Availability

The WT genome is available at GenBank with accession number GCA_036346035.1; the BR genome is available at CNGDC with accession number GWHAOPW00000000; re-sequencing data of 10 WT individuals are available at NCBI SRA with accession number SRP433016; re-sequencing data of 41 BR individuals are available at CNGDC with project number PRJCA003284; re-sequencing data of 12 BL individuals are available at CNGDC with project number PRJCA003284 and at NCBI SRA with accession number SRP471294; RNA-seq data from 20 tissues of the WT individual are available at NCBI SRA with accession number SRP432961; RNA-seq data from the BR individual’s blood, developing primaries and developing tail feathers tissues are available at CNGDC with project number PRJCA003284. Our gene annotation results of the WT and BR genomes are available at FigShare (https://doi.org/10.6084/m9.figshare.24438295.v1).

## Code Availability

All gene annotation code and the corresponding pipeline description are available at https://github.com/zhengchangsulab/A-genome-assebmly-and-annotation-pipeline.

## Author Contributions

CG, JJ, ZS and TD supervised and conceived the project; KW, XG and DW collected tissue samples and conducted molecular biology experiments; SW and SY assembled and annotated the genomes; SW and ZS performed data analysis; and SW and ZS wrote the manuscript.

## Competing interests

The authors declare that they have no competing interests.

## Funding

This work was supported by the National Natural Science Foundation of China (U2002205 and U1702232), Yunling Scholar Training Program of Yunnan Province (2014NO48), Yunling Industry and Technology Leading Talent Training Program of Yunnan Province (YNWR-CYJS-2015-027), Natural Science Foundation of Yunnan Province (2019IC008 and 2016ZA008), and Department of Bioinformatics and Genomics of the University of North Carolina at Charlotte.

## Notes

### Competing Interest Statement

The authors have declared no competing interest.

### Summary of Updates

This study is based in part upon SW's dissertation.

