## Supplementary Note for "Comparative and population genomics analyses of eared pheasants inhabiting highly varying altitudes"

**Primers for RT-PCR validation of 17 new genes**

**1 XR_004170009.1**

|  | Sequence (5'->3') | Tem. strand | Len | Start | End | Tm | GC% | Self com. | Self 3' com. |
| --- | --- | --- | --- | --- | --- | --- | --- | --- | --- |
| Forward primer | GAAACTCCCAGGCTCATCCC | Plus | 20 | 4324 | 4343 | 60.11 | 60.00 | 3.00 | 1.00 |
| Reverse primer | GTGAGTGTTGGACTGAGGCA | Minus | 20 | 4503 | 4484 | 59.89 | 55.00 | 3.00 | 1.00 |
| Product length | 180 | | | | | | | | |

**2 OW982300.1-1**

|  | Sequence (5'->3') | Tem. strand | Len | Start | End | Tm | GC% | Self com. | Self 3' com. |
| --- | --- | --- | --- | --- | --- | --- | --- | --- | --- |
| Forward primer | AATGCGTTTGTGAATGTGGCT | Plus | 21 | 43 | 63 | 59.66 | 42.86 | 4.00 | 0.00 |
| Reverse primer | GTGACTTTTCTCCTCACCACCT | Minus | 22 | 167 | 146 | 59.90 | 50.00 | 4.00 | 0.00 |
| Product length | 125 | | | | | | | | |

**3 OW982314.1-4**

|  | Sequence (5'->3') | Tem. strand | Len | Start | End | Tm | GC% | Self com. | Self 3' com. |
| --- | --- | --- | --- | --- | --- | --- | --- | --- | --- |
| Forward primer | AGTGACTCCGTTTCTCGGGT | Plus | 20 | 388 | 407 | 60.83 | 55.00 | 5.00 | 1.00 |
| Reverse primer | TGCAGAAATAGCAGCCCACC | Minus | 20 | 503 | 484 | 60.68 | 55.00 | 4.00 | 0.00 |
| Product length | 116 | | | | | | | | |

**4 OW982293.1-23**

|  | Sequence (5'->3') | Tem. strand | Len | Start | End | Tm | GC% | Self com. | Self 3' com. |
| --- | --- | --- | --- | --- | --- | --- | --- | --- | --- |
| Forward primer | GGCCACTCAGGCCTTACTTA | Plus | 20 | 130 | 149 | 59.09 | 55.00 | 6.00 | 2.00 |
| Reverse primer | GTTCTGGCTGCATGGTTTCC | Minus | 20 | 232 | 213 | 59.76 | 55.00 | 4.00 | 1.00 |
| **Product length** | 103 | | | | | | | | |

**5 OW982298.1-12**

|  | Sequence (5'->3') | Tem. strand | Len | Start | End | Tm | GC% | Self com. | Self 3' com. |
| --- | --- | --- | --- | --- | --- | --- | --- | --- | --- |
| Forward primer | CAGCTCCCGTTTGGGTATCG | Plus | 20 | 53 | 72 | 60.81 | 60.00 | 4.00 | 2.00 |
| Reverse primer | GGACCCGTTGGAGCACATAC | Minus | 20 | 224 | 205 | 60.74 | 60.00 | 3.00 | 2.00 |
| Product length | 172 |  |  |  |  |  |  |  |  |

**6 XM_048930375.1**

|  | Sequence (5'->3') | Tem. strand | Len | Start | End | Tm | GC% | Self com. | Self 3' com. |
| --- | --- | --- | --- | --- | --- | --- | --- | --- | --- |
| Forward primer | CCGATCTGGGGCAAAGACAT | Plus | 20 | 1477 | 1496 | 60.11 | 55.00 | 4.00 | 2.00 |
| Reverse primer | TCAATCCATCGAACTCGGGC | Minus | 20 | 1622 | 1603 | 60.18 | 55.00 | 4.00 | 2.00 |
| Product length | 146 | | | | | | | | |

**7 OW982301.1-11**

|  | Sequence (5'->3') | Tem. strand | Len | Start | End | Tm | GC% | Self com. | Self 3' com. |
| --- | --- | --- | --- | --- | --- | --- | --- | --- | --- |
| Forward primer | ACGGTAGATCTGCGGTGGAT | Plus | 20 | 16 | 35 | 60.76 | 55.00 | 8.00 | 3.00 |
| Reverse primer | GAAATCCTCTTGCCCCGGTT | Minus | 20 | 148 | 129 | 60.32 | 55.00 | 4.00 | 0.00 |
| Product length | 133 | | | | | | | | |

**8 XM_031605884.1**

|  | Sequence (5'->3') | Tem. strand | Len | Start | End | Tm | GC% | Self com. | Self 3' com. |
| --- | --- | --- | --- | --- | --- | --- | --- | --- | --- |
| Forward primer | CCTGTGCCAGCCTGTAAACT | Plus | 20 | 13228 | 13247 | 60.25 | 55.00 | 5.00 | 1.00 |
| Reverse primer | TCAGATAGCGTTGCGCTGTTA | Minus | 21 | 13477 | 13457 | 60.14 | 47.62 | 5.00 | 2.00 |
| Product length | 250 | | | | | | | | |

**9 XM_048951707.1**

|  | Sequence (5'->3') | Tem. strand | Len | Start | End | Tm | GC% | Self com. | Self 3' com. |
| --- | --- | --- | --- | --- | --- | --- | --- | --- | --- |
| Forward primer | AAGACGATTACAGGGCACCG | Plus | 20 | 1272 | 1291 | 60.11 | 55.00 | 2.00 | 2.00 |
| Reverse primer | GATGCTTCTTGTGAGCGTGC | Minus | 20 | 1487 | 1468 | 60.18 | 55.00 | 4.00 | 2.00 |
| Product length | 216 | | | | | | | | |

**10 XM_040653021.1**

|  | Sequence (5'->3') | Tem. strand | Len | Start | End | Tm | GC% | Self com. | Self 3' com. |
| --- | --- | --- | --- | --- | --- | --- | --- | --- | --- |
| Forward primer | GCTGGGAGCTTGTAAGGAAGT | Plus | 21 | 1344 | 1364 | 60.00 | 52.38 | 4.00 | 1.00 |
| Reverse primer | GCAGCAGTGAGGAGATCGAAA | Minus | 21 | 1539 | 1519 | 60.40 | 52.38 | 4.00 | 0.00 |
| Product length | 196 | | | | | | | | |

**11 XM_048928842.1**

|  | Sequence (5'->3') | Tem. strand | Len | Start | End | Tm | GC% | Self com. | Self 3' com. |
| --- | --- | --- | --- | --- | --- | --- | --- | --- | --- |
| Forward primer | GCAGGAACTGGGGTTACAGA | Plus | 20 | 1378 | 1397 | 59.31 | 55.00 | 3.00 | 0.00 |
| Reverse primer | CACTCATTCCTGAGTTGGGTTC | Minus | 22 | 1618 | 1597 | 58.92 | 50.00 | 6.00 | 0.00 |
| Product length | 241 | | | | | | | | |

**12 XM_042875172.1**

|  | Sequence (5'->3') | Tem. strand | Len | Start | End | Tm | GC% | Self com. | Self 3' com. |
| --- | --- | --- | --- | --- | --- | --- | --- | --- | --- |
| Forward primer | ACACTGGACTGGTATGCAAGG | Plus | 21 | 42 | 62 | 60.00 | 52.38 | 4.00 | 2.00 |
| Reverse primer | GGAGGATCGGGGAGATTCCA | Minus | 20 | 142 | 123 | 60.47 | 60.00 | 4.00 | 2.00 |
| Product length | 101 | | | | | | | | |

**13 novel1**

|  | Sequence (5'->3') | Tem. strand | Len | Start | End | Tm | GC% | Self com. | Self 3' com. |
| --- | --- | --- | --- | --- | --- | --- | --- | --- | --- |
| Forward primer | TTCTGACCTCCTCCGTCCTA | Plus | 20 | 381 | 400 | 59.00 | 55.00 | 3.00 | 2.00 |
| Reverse primer | CGAAAGGAGGCTGTAGCAAG | Minus | 20 | 481 | 462 | 58.64 | 55.00 | 4.00 | 0.00 |
| Product length | 101 | | | | | | | | |

**14 novel9**

|  | Sequence (5'->3') | Tem. strand | Len | Start | End | Tm | GC% | Self com. | Self 3' com. |
| --- | --- | --- | --- | --- | --- | --- | --- | --- | --- |
| Forward primer | GTCCAGAGGTGAAAGTCCAAGT | Plus | 22 | 98 | 119 | 59.90 | 50.00 | 2.00 | 1.00 |
| Reverse primer | GTGCCATCACAGTCACTCCT | Minus | 20 | 218 | 199 | 59.68 | 55.00 | 5.00 | 1.00 |
| Product length | 121 | | | | | | | | |

**15 novel22**

|  | Sequence (5'->3') | Tem. strand | Len | Start | End | Tm | GC% | Self com. | Self 3' com. |
| --- | --- | --- | --- | --- | --- | --- | --- | --- | --- |
| Forward primer | GCGCCTTTATGGGATCCTG | Plus | 19 | 5 | 23 | 58.37 | 57.89 | 6.00 | 2.00 |
| Reverse primer | CATCGCCTTCCATGGGGATTT | Minus | 21 | 239 | 219 | 60.76 | 52.38 | 6.00 | 2.00 |
| Product length | 235 | | | | | | | | |

**16 novel47**

|  | Sequence (5'->3') | Tem. strand | Len | Start | End | Tm | GC% | Self com. | Self 3' com. |
| --- | --- | --- | --- | --- | --- | --- | --- | --- | --- |
| Forward primer | GGAAGAGCATCTCCTTGCCC | Plus | 20 | 94 | 113 | 60.47 | 60.00 | 4.00 | 2.00 |
| Reverse primer | TGGCTGATTCCATGGGAAGAC | Minus | 21 | 195 | 175 | 60.06 | 52.38 | 6.00 | 2.00 |
| Product length | 102 | | | | | | | | |

**17 novel55**

|  | Sequence (5'->3') | Tem. strand | Len | Start | End | Tm | GC% | Self com. | Self 3' com. |
| --- | --- | --- | --- | --- | --- | --- | --- | --- | --- |
| Forward primer | GCAGCTGATGGTGTTTTCCTG | Plus | 21 | 270 | 290 | 60.07 | 52.38 | 6.00 | 1.00 |
| Reverse primer | AGGAGAGGCACACAGTTTTGG | Minus | 21 | 388 | 368 | 60.48 | 52.38 | 3.00 | 0.00 |
| Product length | 119 | | | | | | | | |
